# Exendin-4 disrupts responding to reward predictive incentive cues in rats

**DOI:** 10.1101/315705

**Authors:** Ken T. Wakabayashi, Ajay N. Baindur, Malte Feja, Karie Chen, Kimberly Bernosky-Smith, Caroline E. Bass

**Affiliations:** Department of Pharmacology and Toxicology, Jacobs School of Medicine and Biomedical Sciences, University at Buffalo, State University of New York, Buffalo, NY 14203, USAs; Research Institute on Addictions, University at Buffalo, State University of New York, Buffalo, NY 14203, USA; D’Youville College, Department of Biology and Mathematics, Buffalo, NY, 14201, USA.

## Abstract

Exendin-4 (EX4) is a GLP-1 receptor agonist used clinically to control glycemia in Type-2 Diabetes Mellitus (T2DM), with the additional effect of promoting weight loss. The weight loss seen with EX4 is attributable to the varied peripheral and central effects of GLP-1, with contributions from the mesolimbic dopamine pathway that are implicated in cue-induced reward seeking. GLP-1 receptor agonists reduce preference for palatable foods (i.e. sweet and fat) as well as the motivation to obtain and consume these foods. Accumulating evidence suggest that GLP-1 receptor activity can attenuate cue-induced reward seeking behaviors. In the present study, we tested the effects of EX4 (0.6, 1.2, and 2.4 µg/kg i.p.) on incentive cue (IC) responding. This rat model required rats to emit a nosepoke response during an intermittent audiovisual cue to obtain a sucrose reward (10% solution). EX4 dose-dependently attenuated responding to reward predictive cues, and increased latencies of the cue response and reward cup entry to consume the sucrose reward. Moreover, EX4 dose-dependently decreased the number of nosepokes relative to the number of cue presentations during the session. There was no drug effect on the number of reward cup entries per reward earned during the session, a related reward-seeking behavior with similar locomotor demand. Interestingly, there was a dose-dependent effect of time on the responding to reward predictive cues and nosepoke response latency, such that 2.4 µg/kg of EX4 delayed responding to the initial IC of the behavioral session. Together, these findings suggest that agonism of the GLP-1 receptor with EX4 modulates the incentive properties of cues attributed with motivational significance.

## 1. Introduction

Exendin-4 (EX4) is a glucagon-like peptide-1 (GLP-1) receptor agonist that has been approved to treat type 2 diabetes mellitus (T2DM). EX4 regulates glycemia by activation of peripheral GLP-1 receptors (GLP-1R) in the intestinal tract, driving GLP-1 stimulation of insulin release and increasing gastric emptying time. Central GLP-1 releasing neurons in the nucleus tractus solitarius (NTS) of the dorsal medulla respond to afferent vagal stimuli involving feeding and project widely throughout the brain to modulate satiety (Grill et al., 2007; Holst, 2007; Merchenthaler et al., 1999). In addition to projections to areas involved in homeostatic feeding, some NTS GLP-1 neurons innervate mesolimbic areas involved in regulating motivated reward-seeking behavior (Alhadeff et al., 2012). For example, systemic, intraventricular (ICV), and targeted EX4 microinfusions into the mesolimbic structures including the nucleus accumbens (NAc) and ventral tegmental area (VTA) decreased lever pressing for food rewards and decreased food intake when highly palatable foods and standard chow were available concurrently (Dickson et al., 2012; Yang et al., 2014). Recently, we demonstrated that EX4 dose-dependently attenuated operant responding for a sweetened fat reward under both the fixed ratio 1 and progressive ratio schedules (FR1 and PR, respectively (Bernosky-Smith et al., 2016), with EX4 being more effective in reducing FR1 responding.

It is becoming increasingly clear that GLP-1R agonists can also impact drug-seeking behaviors. EX4 decreases both amphetamine-induced accumbal dopamine release and conditioned place preference (Egecioglu et al., 2013). ICV delivery of EX4 suppresses cocaine-induced phasic dopamine release in the NAc core, but not shell, with no effect on electrically stimulated release (Fortin and Roitman, 2017). Recently, systemic delivery of EX4 blocked cue-induced reinstatement of cocaine-seeking behavior at doses that did not impact measures of food intake, an effect attenuated by delivery of an intra-VTA GLP-1 antagonist (Hernandez et al., 2018). Thus, in addition to the primary effects of EX4 and other GLP-1R agonists on the homeostatic drive for food reinforcers, there is evidence that these drugs can also alter responding to reward associated cues themselves. Further supporting this hypothesis, in diabetic humans GLP-1R agonists have been shown to decrease neuronal activation induced by food cues (e.g. pictures of palatable food) (Farr et al., 2016; Ten Kulve et al., 2016). In rodent models, central administration of EX4 attenuates cue-induced reinstatement of sucrose seeking and conditioned place preference for a high-fat food (Ong et al., 2017).

Pursuing a food or drug reward is a multifaceted process and employs multiple reward-related systems including those that process cues that have been repeatedly paired with reward (Fields et al., 2007; Kelley and Berridge, 2002; Meyer et al., 2016). To test the hypothesis that GLP-1R agonism disrupts the incentive motivational properties (“incentive salience” (Berridge and Robinson, 1998)) attributed to a reward predictive cue, we have explored how EX4 impacts operant responding in a task heavily dependent on incentive cues (ICs). During this task, rats must nosepoke during a random, 8 second intermittent audiovisual cue to receive a sucrose reward. Behavioral metrics indicative of the strength of the IC and primary reinforcer can then be measured. We predicted that EX4 would dose-dependently decrease responding to the number of ICs and increase the latency to respond to ICs and to enter the reward receptacle, indicating that the motivation for the IC and sucrose would both be attenuated.

## 2. Methods

### 2.1 Subjects

Seven male Long Evans rats weighing between ~280-300 grams were purchased from Envigo (Indianapolis, IN) and individually housed under a 12:12 light-dark cycle, with lights on at 3:00 PM and off at 3:00 AM. Food and water were available *ad libitum*. Rats were weighed on a weekly basis. All procedures were reviewed and approved by the University at Buffalo Institutional Animal Care and Use Committee.

### 2.2 Reagents

EX4 (American Peptide Company, Sunnyvale, CA) was suspended in 0.9% saline and stored at −80°C until the day of the experiment. EX4 stocks were thawed on ice and 0.6, 1.2 and 2.4 µg/kg solutions were made. Sucrose was diluted with water to 10% concentration and stored at 4°C.

### 2.3 Behavioral Apparatus

Operant chambers (Med-Associates, Georgia, VT) were housed in sound attenuation cubicles. Each chamber was equipped with illuminated nosepokes located on the left and right of a central liquid receptacle with an infrared entry detector. The liquid cup was filled with an 18-g cannula attached to the bottom of the cup connected to a 10 ml syringe containing 10% sucrose in a syringe pump. A white house light and speaker were located on the wall opposite the nosepokes and reward cup.

### 2.4 IC Operant Task Training and Testing

Rats were acclimated to the testing room for at least 30 min prior to being placed in an operant chamber. Each session lasted 1 hour. The IC task is modified from other procedures described elsewhere (Ambroggi et al., 2011; Wakabayashi et al., 2018; Wakabayashi et al., 2004; Yun et al., 2004). Briefly, rats (n=7) were first trained to nosepoke into the active nosepoke port to receive ~60 µl of 10% sucrose. Initially, a tone and light combination (consisting of an intermittent 2.9 kHz, ~80 dB tone with a 25/20 ms tone-on/off pulse, illumination of the active port while the houselight was turned off) was present during the entire session, except when the rat entered the port. Together, this combination of sound plus light stimuli comprised the IC. When the rat entered the active port, the audiovisual cue was terminated and the syringe pump was activated for 4s, the houselight was illuminated and the tone changed from intermittent to constant; this conditioned stimulus (CS) was distinct from the IC. There was no time-out period and the session terminated when the rat achieved 130 rewards or 1-hr had elapsed. When the rats received at least 100 rewards for two consecutive days, the IC was decreased to 30-s total, and presented on a variable interval 30-s schedule in which inter-trial intervals between IC presentations were randomly selected from a Gaussian distribution (lower and upper limits 15s and 45s, respectively). Entry into the active port during the IC terminated it, followed by reward delivery and presentation of the CS. Nosepokes into the port during the inter-trial interval, or during reward delivery, as well as entries into the inactive nosepoke, were recorded but had no consequences. Once rats achieved responding to 80% or more ICs during two consecutive sessions, the total IC length was set to 8s, with ~100 trials during the 1-hr session. Behavioral sessions were conducted at 9:00 AM, 5 days a week, and trained to a performance criterion of 80% responding to the ICs presented during the session. Stable acquisition of behavior (responding to 80% of ICs for two consecutive sessions) was required prior to testing of the EX4.

### 2.5 Experimental Design, calculations, and statistical analyses

Once performance criteria on the IC task was met, rats were challenged with EX4 in a Latin square design. Rats were administered either saline, 0.6, 1.2 or 2.4 µg/kg of EX4, intraperitoneally (i.p.) 30 min prior to the start of the session. At least three sessions with no pretreatment were allowed before the next challenge dose of EX4. The metrics collected included the: response ratio and latencies to nosepoke into the active port and enter the reward cup, number of nosepokes in the active port for every IC presented, number of cup entries per reward obtained, and which IC was first rewarded. The response ratio equaled the total number of rewarded nosepokes/total number of IC trials presented. For calculations of ED_50_ values, all experimental values were transformed to percent inhibition (% inhibition = 100 × [1 - (test value – mean max value)/ (mean saline value – mean max value)], where 2.4 µg/kg of EX4 provides the maximum value. ED_50_ values were calculated using the least squares linear regression followed by calculation of 95% confidence limits (Bliss, 1967). Data were analyzed using Repeated Measures (RM) one-way ANOVA with a Dunnett’s post-hoc comparison between each dose and the vehicle control condition. To assess drug effects during the session, performance across the three behavioral metrics were compared between the first and last 15 minutes of the session, and statistically analyzed by a RM two-way ANOVA with a Holms-Sidak post-hoc comparison within each dose. To assess if EX4 caused a decrease in motivation for the IC from the onset of the session, or if the rat needed to experience the sucrose reward, we quantified which IC the rats first responded to and analyzed this by a Friedman test.

## 3. Results

EX4 produced a profound dose-dependent decrease in responding to the ICs, along with increases in latency to nosepoke in response to the IC, and latency to enter the reward cup after emitting a rewarded nosepoke (Figure 1). The mean response ratio ± SEM after vehicle was 0.87 ± 0.03, and 0.6, 1.2 and 2.4 µg/kg EX4 produced decreases in the response ratio to 0.79 ± 0.03, 0.61 ± 0.02, and 0.33 ± 0.07, respectively. The response ratio was converted to a % inhibition, and analysis of the dose response curve determined the ED_50_ to be 1.13 (0.96-1.32) µg/kg. The 1.2 and 2.4 µg/kg dose produced a significant increase in % inhibition of the response ratio compared to vehicle (RM one-way ANOVA F_3,18_ = 29.39, p < 0.0001). We also observed increases in the average nosepoke and reward cup latencies. The mean nosepoke latency ± SEM increased from 1.65 ± 0.14 seconds after vehicle to 2.13 ± 0.19, 2.28 ± 0.11, and 2.75 ± 0.08 seconds after 0.6, 1.2 and 2.4 µg/kg EX4, respectively. Likewise the mean latency to enter the reward cup after a rewarded nosepoke was 0.61 ± 0.03 seconds after saline, and increased to 0.71 ± 0.04, 0.824 ± 0.06, and 1.10 ± 0.11 seconds after 0.6, 1.2 and 2.4 µg/kg EX4, respectively. We then converted these two metrics to a percent change in latency, and analyzed the resulting dose response curves. The ED_50_ of the percent change in nosepoke latency was 0.79 (0.51-1.22) µg/kg EX4, and both 1.2 and 2.4 µg/kg EX4 doses significantly differed from vehicle (RM one-way ANOVA analysis F_3,18_ = 10.3, p < 0.0005). Analysis of the dose response curve for the percent change in reward cup latency produced an ED_50_ of 1.09 (0.79-1.5) µg/kg EX4 and RM one-way ANOVA analysis (F_3,18_ = 12.14, p < 0.0005) which determined that all three doses of EX4 significantly increased the latency to enter the reward cup compared to vehicle.

**Figure 1.**
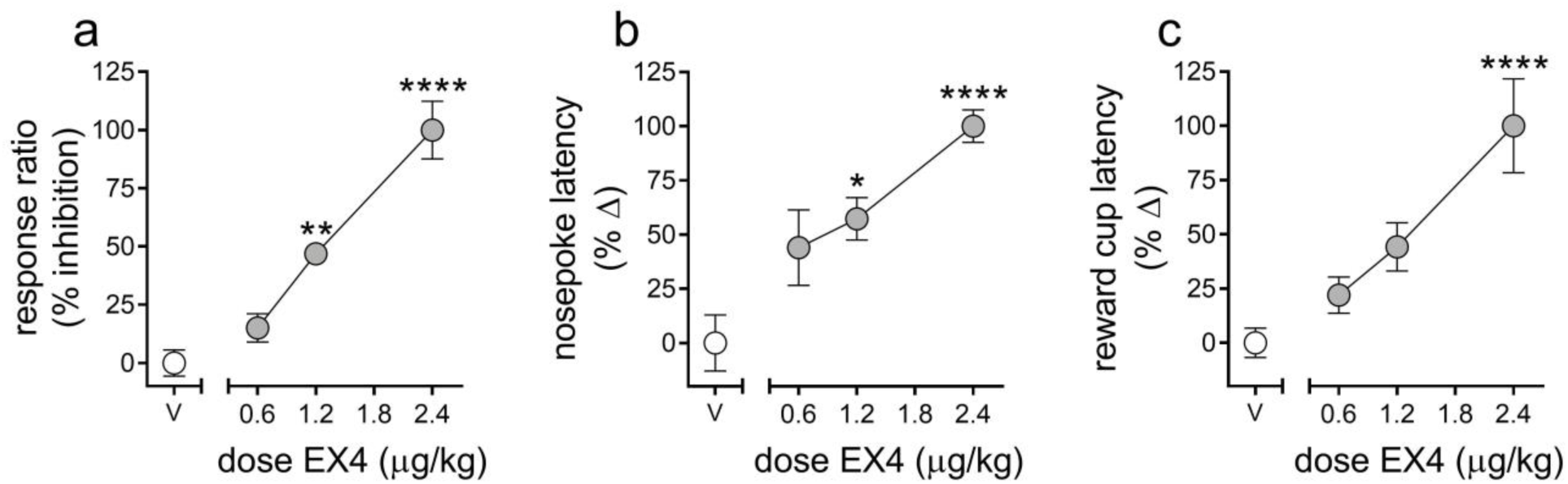
Effects of EX4 on responding during the incentive cue-instrumental task. EX4 dose-dependently increased the % inhibition of the response ratio (a), and the % change in latencies to nosepoke (b) and enter the reward cup (c) to collect the reward. * p < 0.05, ** p < 0.01, **** p < 0.0005 compared to the vehicle treatment value.

Rats can emit additional unrewarded nosepokes in the active port either while the reward is being delivered or between IC presentations, leading to more total active nosepokes under vehicle conditions than total ICs presented during the session. EX4 robustly and dose-dependently decreased nosepokes in each session, with vehicle pretreatment producing a mean of 117.14 ± 7.58, while 0.6, 1.2 and 2.4 µg/kg EX4 decreased the mean total nosepokes to 101.71 ± 6.87, 78 ± 4.85, and 39.86 ± 7.64, respectively (Figure 2a, RM one-way ANOVA F_3,18_ = 22.54, p < 0.0001). This decrease in nosepokes could result from generalized locomotor inhibition. However, the total number of reward cup entries during the session, a behavioral response with a similar locomotor demand as nosepokes, was less impacted by EX4 (Figure 2b, RM one-way ANOVA F_3,18_ = 7.46, p < 0.005). The mean reward cup entries for vehicle were 144 ± 16.83, while 0.6, 1.2 and 2.4 µg/kg EX4 decreased this to 130.4 ± 5.97, 142 ± 18.46, and 71.29 ± 13.01, respectively. However, only the highest dose of EX4, 2.4 µg/kg, produced a significant decrease in reward cup entries compared to vehicle.

**Figure 2:**
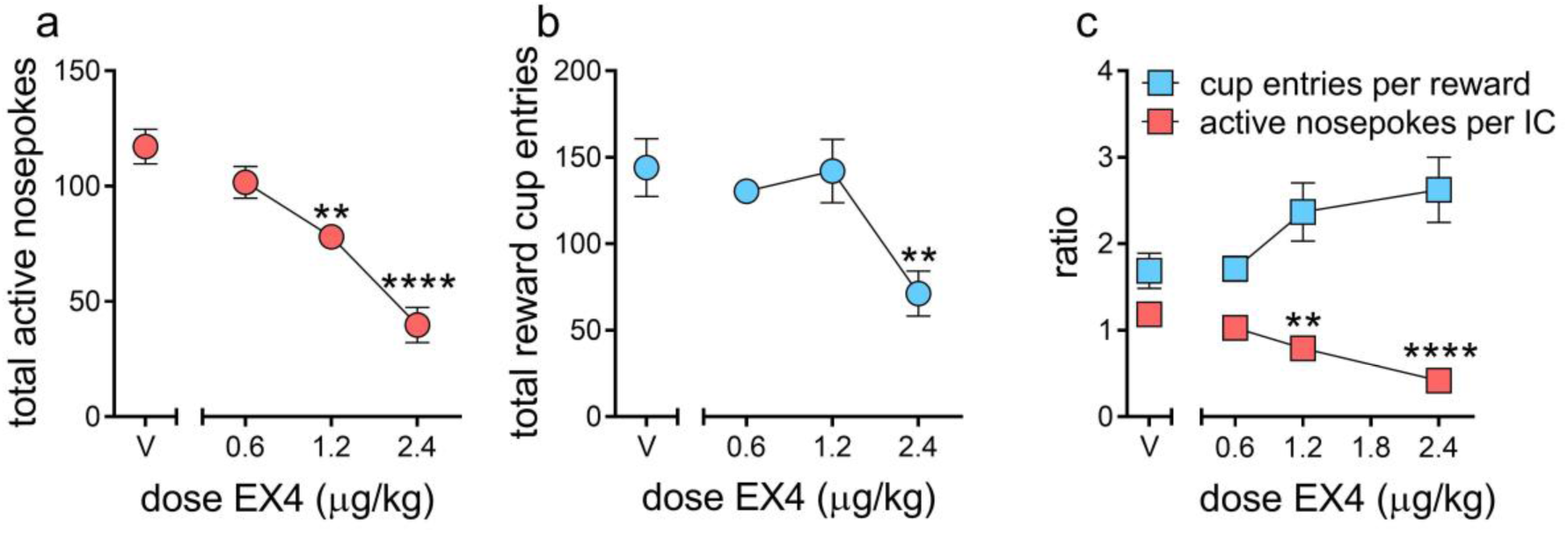
EX4 decreases motivation for the IC while motivation for sucrose is retained. The total number of active nosepokes during the session dose-dependently decreased (a), while the number of total reward cup entries (b) was less attenuated by EX4. Normalizing responses as a ratio of cup entries per reward (c, blue squares) or active nosepokes per IC (red squares) showed that active nosepokes per IC was dose-dependently decreased, while the ratio of reward cup entries per reward earned trended upward. ** p < 0.01, **** p < 0.0005 compared to the vehicle treatment value.

When taking into account the number of opportunities to earn a reward during a session, rats generally emitted 1.18 ± 0.08 nosepokes per IC (vehicle, Figure 2c). EX4 dose-dependently decreased this ratio to 1.03 ± 0.07, 0.79 ± 0.05, and 0.42 ± 0.08 after 0.6, 1.2 and 2.4 µg/kg EX4, respectively, and 1.2 and 2.4 µg/kg EX4 doses significantly decreased this ratio compared to vehicle (RM one-way ANOVA F_3,18_ = 21.47, p < 0.001). This decrease in nosepoke per IC ratio is proportional to the % inhibition in the response ratio (Figure 1). Notably, normalizing the number of reward cup entries to the number of rewards earned during the session revealed that rats entered the reward cup 1.69 ± 0.21 times for each reward earned during the session after vehicle, while 0.6, 1.2 and 2.4 µg/kg EX4 increased this ratio to 1.71 ± 0.15, 2.37 ± 0.34, and 2.62 ± 0.38, respectively. While the reward cup entries per reward increased dose-dependently, this trend did not reach significance (RM one-way ANOVA, (F_3,18_ = 3.07, p=0.0575).

We next determined the effect of EX4 over the course of the 1-hr session by comparing response ratio, nosepoke latency and reward cup latency in the first 15 min of the 1-hr session with the last 15 min of the session. The mean response ratio for vehicle between the first and last 15 minute bins were 0.84 ± 0.04 and 0.84 ± 0.05, while EX4 decreased this to 0.88 ± 0.04 and 0.76 ± 0.05 (0.6 µg/kg), 0.78 ± 0.06 and 0.59 ± 0.06 (1.2 µg/kg), 0.17 ± 0.07 and 0.50 ± 0.10 (2.4 µg/kg) respectively. We determined there was a significant main effect of dose (Figure 3, RM two-way ANOVA; F^3, 18^ = 27.27, p < 0.0001) with a significant dose X session bin interaction (F^3,18^ = 13.32, p < 0.0001), such that a majority of rats decreased their response ratio between the beginning and end of the session after 1.2 µg/kg EX4, but exhibited the opposite pattern after 2.4 µg/kg. The mean latencies to nosepoke for vehicle between the first and last 15 minute bins were 1.39 ± 0.18 and 1.79 ± 0.11 seconds, while EX4 increased this to 1.75 ± 0.23 and 2.26 ± 0.20 (0.6 µg/kg), 2.34 ± 0.21 and 2.37 ± 0.19 (1.2 µg/kg), 3.96 ± 0.86 and 2.24 ± 0.17 seconds (2.4 µg/kg), respectively. Analysis of nosepoke latency in response to the cue demonstrated that rats had a greater latency to nosepoke at the beginning of the session after 2.4 µg/kg (RM two-way ANOVA, main effect of session bin F_3,18_ = 9.203, p < 0.001, session bin X treatment interaction, F_3,18_ = 7.017, p < 0.005), with no other observable differences with lower doses. Conversely, the mean latencies for vehicle between the first and last bins were 0.55 ± 0.04 and 0.68 ± 0.06, while EX4 increased this to 0.71 ± 0.06 and 0.68 ± 0.06 (0.6 µg/kg), 0.83 ± 0.06 and 0.81 ± 0.06 (1.2 µg/kg), 1.07 ± 0.10 and 1.0 ± 0.08 seconds (2.4 µg/kg). There was no overall difference between the beginning and end of the session (RM two-way ANOVA, main effect of dose, F_3,18_ = 13.46, p < 0.0001). Under control conditions, rats normally responded to either the first or second IC presented in the session. This effect increased in several subjects dramatically at the highest dose of EX4 tested, although this effect because of within group variance did not quite reach statistical significance (p = 0.06).

**Figure 3:**
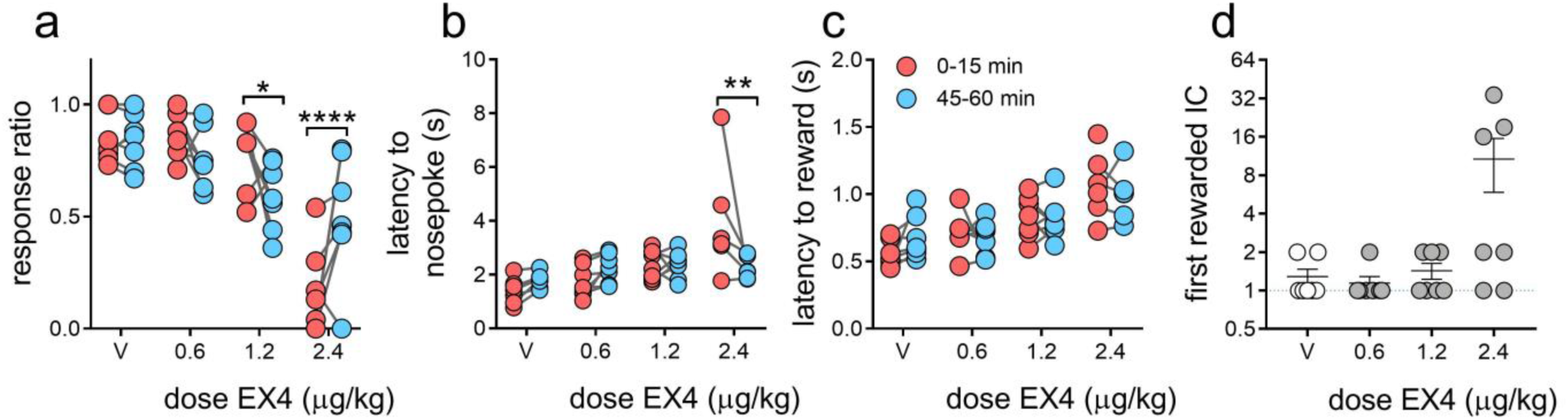
Differential effects of EX4 within an IC session. EX4 had contrasting dose-dependent effects on response ratio (a) during the beginning (red circles) and end (blue circles) of a session. EX4 dramatically increased the latency to nosepoke during the beginning of a session at 2.4 µg/kg EX4 (b), while latency to reward was not impacted within the session for any dose of EX4 (c). After EX4 treatment, a subset of rats responded to a later IC during the session (d). Note that circles represent individual subjects. Some individuals in B and C lack data points since they did not respond during the first or last 15 min of the session. * p < 0.05, ** p < 0.01, **** p < 0.0005 comparing the first and last 15 minutes of the session.

## 4. Discussion

In the present study, we determined the effects of EX4 in a novel IC task to establish whether this drug affected both the motivation to obtain food rewards, as well as the ability of ICs to promote reward seeking behaviors. Our data show that EX4 attenuates the incentive salience attributed to reward predictive cues. While previous studies have suggested that GLP-1 signaling contributes to consumption, satiety and the motivation to work for food, it was unclear whether EX4 and other GLP-1R agonists were acting solely via the homeostatic drive for food reward. Here, our findings using the IC task suggest that GLP-1 also directly modulates the motivational properties of cues predicting reward availability. Specifically, EX4 dose-dependently attenuated responding to these ICs, as well as increased the latencies to respond to both the cues and to enter the reward cup to consume the sucrose reward (Figure 1).

NTS neurons producing endogenous GLP-1 projecting to the VTA and NAc may modulate the rewarding value of food and other rewards (Alhadeff et al., 2012; Merchenthaler et al., 1999). Recent studies have shown that systemic and intra-VTA administration of EX4 can attenuate cue-induced cocaine seeking at doses that do not impact food intake (Hernandez et al., 2018). Central administration of EX4 has recently been shown to attenuate cocaine-induced phasic dopamine signaling in the NAc (Fortin and Roitman, 2017), possibly by increasing GABA_A_ receptor mediated inhibition (Farkas et al., 2016; Korol et al., 2015a; Korol et al., 2015b). Our present findings support the hypothesis that GLP-1R agonists impact reward-seeking behaviors by disrupting the motivational salience attributed to reward predictive incentive cues.

Others have suggested that operant reward-seeking performance can be parsed into two components quantifying motivation: *response choice*, which represents making a choice between different possible alternative actions, and *response vigor*, which measures the latency with which the chosen action is performed (Niv et al., 2007). In our behavioral task, the choice to enter the active nosepoke or reward cup are quantifiable elements of response choice, while the latency to nosepoke in response to an IC, and enter the reward cup after a reward has been delivered, are indices of response vigor. We found that behaviors more tightly associated with responding to the IC, such as nosepoking, are robustly attenuated by EX4 (Figure 2a). Likewise, reward centered behaviors (i.e. the number of reward cup entries) are also attenuated, though only at the highest 2.4 µg/kg dose (Figure 2b). When nosepokes and reward cup entries were normalized to their corresponding events during the session (e.g. the number of ICs or rewards delivered, respectively), it appears that the motivation for the cues (number of active nosepokes per IC) were dose-dependently attenuated, while motivation for the reward increased (i.e. number of reward cup entries per reward, Figure 2c). Although this increase was not significant (p = 0.06), nonetheless EX4 did not decrease this metric at any dose. Together, these data suggest that in this IC task, increasing doses of EX4 selectively impaired motivation for the IC, including both response choice and vigor, while motivated behavior associated with the reinforcer itself was largely retained.

The most parsimonious explanation for attenuation of responding in our IC task is that EX4 inhibited locomotor activity or reduced appetite via an increase in nausea, two well-known side effects of GLP-1R agonists. However, the rats performed the IC task at the same level as vehicle at the 0.6 dose, and to some extent the 1.2 µg/kg EX4 dose, even though these doses have been previously reported to decrease locomotor behavior (Bernosky-Smith et al., 2016). Further, while EX4 inhibited the total number of nosepokes at the 1.2 and 2.4 µg/kg doses, only the highest dose, 2.4 µg/kg, inhibited reward cup entries (Figure 2), even though both metrics require a similar degree of motor activity. Specific to nausea, we have previously reported that EX4 did not induce pica in a kaolin test at any of these doses (Bernosky-Smith et al., 2016). It is possible that the IC task is more sensitive to visceral malaise than the kaolin test, and therefore the decrease in IC responding, particularly at higher doses of EX4 could have a nausea component. However, rats entered the reward cup much more frequently per reward at doses that decreased nosepokes in the active port (1.2 and 2.4 µg/kg, Figure 2c). This implies that nausea is not a major contributor to performance in the IC task, as sucrose seeking was still intact to some extent. Nonetheless future experiments should be performed to determine the sensitivity of the IC task to other drugs known to induce nausea.

Another intriguing finding from this study is that EX4 had differential effects depending on the time of the session and the dose administered. We did not observe any changes in response ratio, nosepoke latency or reward latency at vehicle or the 0.6 µg/kg dose (Figure 3). However, at 1.2 µg/kg, while the response ratio was unchanged during the first 15 minutes, it decreased significantly in the last 15 minutes of the 1-hr session. Yet, at the 2.4 µg/kg dose, the rats had a suppressed response ratio at the beginning of the session, but most recovered by the end and were performing at vehicle levels. This interaction was absent in the nosepoke latency, where the rats were unchanged between the beginning and the end of the session at vehicle, 0.6 and 1.2 µg/kg doses. However, they exhibited significantly higher latencies at the beginning of the session after 2.4 µg/kg, but recovered completely by the end of the session, again where they performed at vehicle levels. The latency to enter the reward cup was dose-dependently increased by EX4, but there were no significant differences between the beginning and end of the session. Together these data indicate that the 1.2 µg/kg dose of EX4 decreased responding to the IC over time during the session, perhaps indicating a decreased satiety threshold, while the higher dose of 2.4 µg/kg was characterized by a delay in starting to respond to the IC, which likely involves different mechanisms, potentially nausea. However, even in this case, when the rats did respond to the IC, their reward centered vigor (i.e. latency to enter the reward cup) remained unchanged during the session.

Further, it has been proposed that changes in incentive motivational processes should occur at the onset of the session, independent of exposure to the reward (Berridge, 2012). In our study, the rats demonstrated a dose-dependent EX4 induced decrease in motivation for the cue from the onset of the session, as indicated by a decrease in response ratio and nosepoke latency during the first 15 minutes of the session, and a delay in the first IC they responded to in the task (Figure 3d), indicating that the 2.4 µg/kg dose of EX4 impacts primarily the motivational salience attributed to the cue. However, it is important to note that we also observed substantial individual variation in the latency to nosepoke and first IC rewarded at the 2.4 µg/kg dose. This is similar to results in our previous study in which 2.4 µg/kg EX4 increased the latency to the first lever press in a model of sweetened fat self-administration (Bernosky-Smith et al., 2016). Such individual differences should be explored further in future studies of EX4 on motivation. Moreover, it should be noted that EX4 is long-acting and should be pharmacologically active well past the 1-hr session, ruling out a return to baseline due to pharmacokinetic factors.

In summary, we found that systemic EX4 preferentially attenuates responding to cues attributed with incentive salience. These behaviors are dissociable from other components of reward seeking using a dynamic and robust free-operant model that is heavily reliant on ICs and can quantify elements of response choice and response vigor. Future studies will need to determine whether GLP-1R regulation of mesolimbic circuits is the primary mechanism by which EX4 decreases the incentive salience of reward cues.

## Acknowledgements

This research was supported by the State University of New York BRAIN Network of Excellence Postdoctoral Fellow program (K.T.W), the Whitehall Foundation 2017-12-98 (C.E.B.), the National Institutes of Health T32 AA007583 (K.T.W), and R21 DA043190 (C.E.B). We thank Dr. Derek Daniels for his helpful advice.

